# Specificity and selective advantage of an exclusion system in the integrative and conjugative element ICE*Bs1* of *Bacillus subtilis*

**DOI:** 10.1101/2021.01.25.428195

**Authors:** Kathleen P. Davis, Alan D. Grossman

**Affiliations:** Department of Biology, Massachusetts Institute of Technology, Cambridge, MA 02139

## Abstract

Integrative and conjugative elements (ICEs) are mobile genetic elements capable of transferring their own and other DNA. They contribute to the spread of antibiotic resistances and other important traits for bacterial evolution. Exclusion is a mechanism used by many conjugative plasmids and a few ICEs to prevent their host cell from acquiring a second copy of the cognate element. ICE*Bs1* of *Bacillus subtilis* has an exclusion mechanism whereby the exclusion protein YddJ in a potential recipient inhibits the activity of the ICE*Bs1-*encoded conjugation machinery in a potential donor. The target of YddJ-mediated exclusion is the conjugation protein ConG (a VirB6 homolog). Here we defined the regions of YddJ and ConG that confer exclusion specificity and determined the importance of exclusion to host cells. Using chimeras that had parts of ConG from ICE*Bs1* and the closely related ICE*Bat1* we identified a putative extracellular loop of ConG that conferred specificity for exclusion by the cognate YddJ. Using chimeras of YddJ from ICE*Bs1* and ICE*Bat1* we identified two regions in YddJ needed for exclusion specificity. We also found that YddJ-mediated exclusion reduced death of donor cells following conjugation into recipients. Donor death was dependent on the ability of transconjugants to themselves become donors and was reduced under osmo-protective conditions, indicating that death was likely due to alterations in the donor cell envelope caused by excessive conjugation. We postulate that elements that can have high frequencies of transfer likely evolved exclusion mechanisms to protect the host cells from excessive death.

**Importance:** Horizontal gene transfer is a driving force in bacterial evolution, responsible for the spread of many traits, including antibiotic and heavy metal resistances. Conjugation, one type of horizontal gene transfer, involves DNA transfer from donor to recipient cells through conjugation machinery and direct cell-cell contact. Exclusion mechanisms allow conjugative elements to prevent their host from acquiring additional copies of the element, and are highly specific enabling hosts to acquire heterologous elements. We defined regions of the exclusion protein and its target in the conjugation machinery that convey high specificity of exclusion. We found that exclusion protects donors from cell death during periods of high transfer. This is likely important for the element to enter new populations of cells.

## Introduction

Integrative and conjugative elements (ICEs, also called conjugative transposons) play a major role in bacterial evolution by contributing to the spread of genetic material, including genes for antibiotic resistances, pathogenesis, symbiosis, and metabolic functions (1–3). ICEs are typically found integrated into the host chromosome. Under certain conditions they can excise and transfer to a new host through conjugation machinery encoded by the element (4,5), thus enabling their spread through a population of bacterial cells. The conjugation machinery encoded by most ICEs is a type 4 secretion system (T4SS) (1) and the genes that confer various phenotypes to the host cells are typically not required for conjugation and are called cargo genes. The conjugation machineries from many ICEs are also capable of transferring (mobilizing) other elements, notably plasmids, to new host cells, allowing for dissemination of elements that do not encode their own conjugation machinery (6–8).

ICE*Bs1* is relatively small (~20 kb) and present in a unique site (in *trnS-leu2*) in most strains of *Bacillus subtilis* (9,10). DNA damage to its host cell, or crowding by *B. subtilis* cells that do not contain ICE*Bs1* both lead to de-repression of transcription of ICE*Bs1* genes and subsequent excision and potential transfer of the element. ICE*Bs1* can be activated in >90% of cells in a population by overproduction of the element-encoded activator protein RapI, making the element readily amenable to population-based studies (9,11,12). ICE*Bs1* has three known mechanisms for inhibiting its host cell from receiving an additional copy of element: 1) inhibition of ICE*Bs1* activation by cell-cell signaling from neighboring cells that already contain a copy of the element (9); 2) repressor-mediated immunity (13); and 3) exclusion (12). Exclusion is a key part of conjugative plasmid biology and most conjugative plasmids appear to have an exclusion system (14). In the F-plasmid of *E. coli*, exclusion protects host cells against lethal zygosis, a phenonemon in which host cells that serve as recipients during excessive transfer events die, likely due to cell wall damage (15–18). In addition, exclusion prevents cells from having recombination events that result in deletions and defective plasmid copies (19–21).

In general, exclusion systems are mediated by a single protein encoded by the element, that is localized to the membrane of the host cell, where it is in position to inhibit cognate conjugation machinery (14). Identified exclusion proteins tend to be fairly small, and membrane attachment is in the form of one or more transmembrane domains, or lipid modification, or both. The target protein in the donor has been identified for exclusion system systems from the F/R100 family of plasmids (22,23), the R64/R62Ia plasmids (24), and the SXT/R391 ICEs (25,26) and ICE*Bs1* (12). ICE*Bs1* is the only ICE from Gram-positive bacteria that is known to have exclusion system.

In ICE*Bs1*, the element-encoded exclusion protein YddJ specifically inhibits its cognate conjugation machinery by targeting the conjugation protein ConG in would-be donor cells, thereby inhibiting transfer of DNA into a cell that already contains ICE*Bs1* (12). ConG, a homolog of VirB6 in the pTI conjugation system from *A. tumefaciens*, is a membrane protein with seven predicted transmembrane segments, and essential for function of the ICE*Bs1* conjugation system (27). Exclusion protects the viability of ICE*Bs1* host cells under conditions that promote conjugation, although it was not clear whether ICE*Bs1* donors, recipients, or both were being protected (12).

Here, we identify the regions in YddJ and ConG that determine the specificity of exclusion. We found that exclusion promotes viability of ICE*Bs1* donor cells by limiting ICE*Bs1* transfer from new transconjugants back into the original donors.

## Results

### Rationale and experimental approach

Exclusion specificity in ICE*Bs1* was established using *conG* and *yddJ* from ICE*Bat1* in place of their homologues in ICE*Bs1* (12). Here, to define the regions of each gene needed to confer specificity, we made chimeras between *conG* or *yddJ* from ICE*Bs1* and ICE*Bat1*. To study the effects of exclusion we used experimental conditions that bypass both cell-cell signaling and immunity. Cell-cell signaling is bypassed by overexpressing *rapI* from an inducible promoter (Pxyl-*rapI*) in ICE-containing donor cells (9). Repressor-mediated immunity in potential recipient cells is bypassed by expressing *yddJ* from an exogenous locus under control of a strong promoter {Pspank(hy)-*yddJ*} in the absence ICE*Bs1* (12).

### Identification of regions of ConG that are essential for exclusion specificity

Regions and resides of ConG and YddJ that are needed for exclusion specificity must be divergent between the proteins from ICE*Bs1* and ICE*Bat1*. There are two main regions of divergence in ConG (12). One includes residues 276-295 of both ConG_Bs1_ and ConG_Bat1_ (Fig. 1), and is predicted to be a loop between the putative third and fourth transmembrane regions. Exclusion-resistant mutations in *conG* are in this loop region (12). The C-terminal region of ConG is also divergent between the two elements. This region is predicted to be a large extracellular domain.

**Figure 1.**
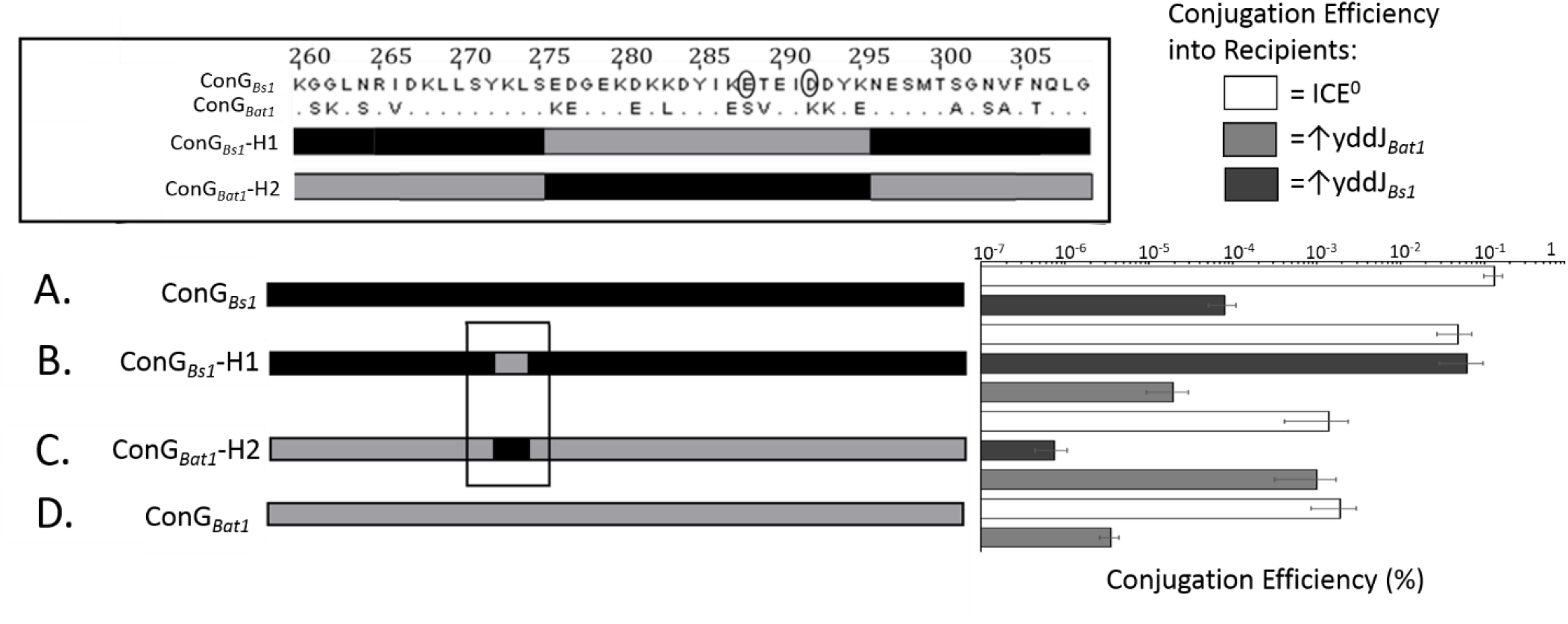
Regions of ConG_Bs1_ and ConG_Bat1_ that confer specificity of exclusion. **Top left**. Comparison of the indicated regions ConG from ICE*Bs1* and ICE*Bat1*. Amino acids that differ between the two proteins in this region are indicated. The two circled residues are sites of mutation that makes ConG insensitive to exclusion (12). The bars below the sequence indicate the regions in the hybrid proteins from ICE*Bs1* (black) and ICE*Bat1* (gray). **Bottom. Left**. Cartoon of ConG present in the donor strains. Regions of ConG from ICE*Bs1* (black) and ICE*Bat1* (gray) are indicated. **A)** ConG_Bs1_ (KPD225); **B)** ConG_Bs1_-H1 (KPD136); **C)** ConG_Bat1_-H2 (KPD135); and **D)** ConG_Bat1_ (KPD224). **Right, top and bottom**. Conjugation efficiencies of the indicated donors (left) into recipients with no YddJ (open, white bars; CAL89); YddJ from ICE*Bs1* (black bars; strain MA982); YddJ from ICE*Bat1* (gray bars; KPD219). Conjugation efficiency is calculated as the CFU/ml of transconjugants divided by the CFU/ml of donors at the start of mating, and is multiplied by 100% to convert to a percentage. Data bars represent averages from three independent experiments, with error bars depicting standard deviations.

We found that amino acids 276-295 in ConG were sufficient to confer specificity. We replaced amino acids 276-295 in ConG_Bs1_ with the corresponding residues from ConG_Bat1_, generating ConG_Bs1_-Bat1(276-295), referred to as ConG_Bs1_-H1 (Fig. 1A, B). This hybrid protein was functional in conjugation with the ICE*Bs1* conjugation machinery. The conjugation efficiency was ~5% (~5 transconjugants per 100 initial donors) into recipients that did not contain YddJ (or ICE*Bs1*). When recipients produced YddJ_Bat1_, the conjugation efficiency was reduced by a factor of ~10^−3^, which we refer to as ~1,000-fold exclusion (Fig. 1B). Exclusion is the ratio of transconjugants into recipients without (no exclusion) versus with *yddJ*. This level of exclusion is similar to that observed when both ConG and YddJ were from ICE*Bat1* (Fig. 1D). In contrast, when recipients produced YddJ_Bs1_, there was no detectable change in the conjugation efficiency giving exclusion of ~1 (no exclusion) (Fig. 1B). Based on these results, we conclude that amino acids 276-295 of ConG from ICE*Bat1* are sufficient to confer exclusion specificity to YddJ from ICE*Bat1*.

We also made the reciprocal replacement, replacing residues 276-295 from ConG_Bat1_ with those from ConG_Bs1_ (Fig. 1A, C). This hybrid ConG_Bat1_-Bs1(276-295), referred to as ConG_Bat1_-H2, was functional in conjugation with the ICE*Bs1* conjugation machinery, but less so than wild type or the other hybrid. The reduced transfer efficiency was expected based on previous analyses substituting ConG_Bat1_ for ConG_Bs1_ in the context of the ICE*Bs1* conjugation machinery (12). The conjugation efficiency was ~0.1% transconjugants per donor into recipients that did not contain YddJ. When recipients produced YddJ from ICE*Bs1*, exclusion was ~1,000-fold, similar that when both ConG and YddJ were from ICE*Bs1* (Fig. 1C). In contrast, when recipients produced YddJ_Bat1_, there was no detectable exclusion (Fig. 1C). Based on these results, we conclude that amino acids 276-295 of ConG from ICE*Bs1* are sufficient to confer exclusion specificity to YddJ from ICE*Bs1*.

Together, the results above indicate that residues 276-295 of both ConG_Bs1_ and ConG_Bat1_ confer specificity of exclusion. These might not be the only residues that contribute to specificity of exclusion, but without these key residues from the cognate ConG, no exclusion by YddJ is observed. With these key residues, exclusion is virtually indistinguishable from that for donors expressing the wild type cognate ConG protein.

### Identification of YddJ Regions Essential for Exclusion Specificity

To identify regions of YddJ_Bs1_ and YddJ_Bat1_ that confer exclusion specificity, we used a similar approach of generating hybrids and testing whether recipient strains expressing these hybrid proteins could exclude ICE*Bs1* using ConG_Bs1_ or ConG_Bat1_. There are two main regions of sequence dissimilarity between the YddJ homologs (Fig. 2A). We made and tested three functional hybrid proteins, focusing on these regions (Fig. 2B).

**Figure 2.**
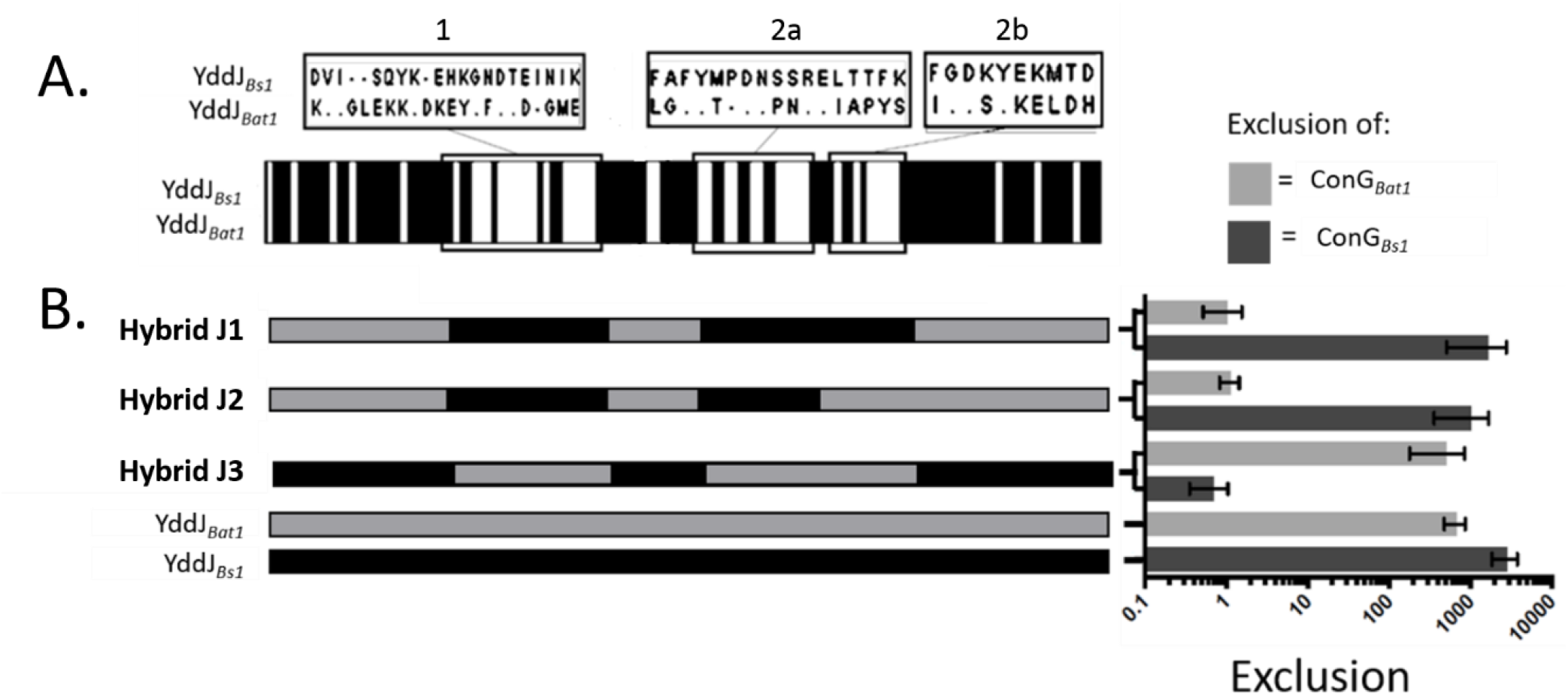
Regions of YddJ_Bs1_ and YddJ_Bat1_ that confer specificity of exclusion. **A)** Protein sequence of the indicated regions of YddJ from ICE*Bs1* and ICE*Bat1*. Amino acids that differ between the two proteins in these regions are indicated. The bars below the sequence compare the proteins across their entire lengths. Black and white indicate sequence identity and divergence, respectively. **B)** Exclusion (right) of the ICE*Bs1* conjugation machinery that contains ConG_Bs1_ (black bars; KPD225) or ConG_Bat1_ (gray bars; KPD224) by the indicated YddJ proteins (left): hybrid J1 (KPD132); hybrid J2 (KPD131); hybrid J3 (KPD128); YddJ_Bat1_ (KPD219); YddJ_Bs1_ (MA982). Exclusion was calculated as conjugation efficiency into a recipient without YddJ (no exclusion; CAL89) divided by that into a recipient expressing the indicated YddJ. Data bars represent averages from three independent experiments, with error bars depicting standard deviations.

#### Hybrid J1, YddJ_Bat1_-Bs1(30-48)(65-81)(86-95)

We made constructs that replaced amino acids from regions 1, 2a and 2b (amino acids 30-50, 67-82, and 87-96) in YddJ_Bat1_ with the corresponding residues (30-48, 65-81, and 86-95, note slightly different numbering) from YddJ_Bs1_ to make hybrid J1, formally known as YddJ_Bat1_-Bs1(30-48)(65-81)(86-95) (Fig. 2B). Hybrid J1 was able to exclude an element that had conjugation machinery with ConG_Bs1_ (exclusion ~1600), but unable to exclude conjugation machinery with ConG_Bat1_ (Fig. 2B). These results indicate that the specificity of YddJ_Bat1_ had been switched to that of YddJ_Bs1_ and that these three regions were sufficient to confer specificity.

#### Hybrid J2, YddJ_Bat1_-Bs1(30-48)(65-81)

We made a construct similar to hybrid J1, but only replaced amino acids in regions 1 and 2a (amino acids 30-50 and 67-82) in YddJ_Bat1_ with the corresponding residues (30-48 and 65-81, respectively) from YddJ_Bs1_ to generate hybrid J2, formally known as YddJ_Bat1_-Bs1(30-48)(65-81) (Fig. 2B). Hybrid J2 excluded a donor with ConG_Bs1_ (exclusion ~1000), but did not exclude a donor with ConG_Bat1_ (Fig. 2B). These results indicate that the specificity of YddJ_Bat1_ had been switched to that of YddJ_Bs1_ and that these two regions were sufficient to change the specificity. They also indicate that amino acids in region 2b (resides 86-95) of YddJ from ICE*Bs1* contributed little if anything to specificity in this context.

#### Hybrid J3, YddJ_Bs1_-Bat1(30-50)(67-82)(87-96)

We made a construct that replaced amino regions 1, 2a and 2b (amino acids 30-50, 67-82, and 87-96) in YddJ from ICE*Bs1* with the corresponding residues from YddJ from ICE*Bat1* to make hybrid J3, formally known as YddJ_Bs1_-Bat1(30-50)(67-82)(87-96) (Fig. 2B). This hybrid is essentially the reciprocal of hybrid J1. Hybrid J3 was able to exclude an element that had conjugation machinery with ConG_Bat1_, but unable to exclude conjugation machinery with ConG_Bs1_ (Fig. 2B). These results indicate that the specificity of YddJ_Bs1_ had been switched to that of YddJ_Bat1_ and that these three regions were sufficient to confer specificity.

We made another hybrid (J4), similar to hybrid J3, but that replaced only regions 1 and 2a (not region 2b) of YddJ_Bs1_ with the corresponding two regions from YddJ_Bat1._ Formally, this hybrid is known as YddJ_Bs1_-Bat1(30-50)(67-82). This hybrid had very little if any exclusion of a donor with ConG_Bat1_ and no detectable exclusion of a donor with Con*G*_Bs1_. These results are largely uninterpretable. It is possible that key residues needed for exclusion were missing. It is also possible that the protein is not functional (perhaps mis-folded). Nonetheless, together, our results indicate that regions 1, 2a, and 2b, and in the context of YddJ_Bat1_, regions 1 and 2a from YddJ_Bs1_ are sufficient to confer exclusion specificity.

### YddJ hybrid proteins can exclude conjugation machinery with ConG hybrid proteins

As a final demonstration of specificity, we tested the ability of hybrid J1 and J3 to exclude conjugation machinery containing the ConG hybrids H1 and H2. Hybrid J1 (YddJ_Bat1_ with regions 1, 2a and 2b from YddJ_Bs1_) was able to inhibit ConG_Bat1_-H2 (ConG_Bat1_ with amino acids 276-295 from ConG_Bs1_), but not ConG_Bat1_ (Fig. 3). Likewise, hybrid J3 (YddJ_Bs1_ with regions 1, 2a and 2b from YddJ_Bat1_) was able to inhibit ConG_Bat1_-H1, but not ConG_Bs1_ (Fig. 3). Together, our results demonstrate that the key residues in ConG_Bs1_ and ConG_Bat1_, and in YddJ_Bs1_ and YddJ_Bat1_, are sufficient to generate the exclusion specificity of their counterpart wild type proteins.

**Figure 3.**
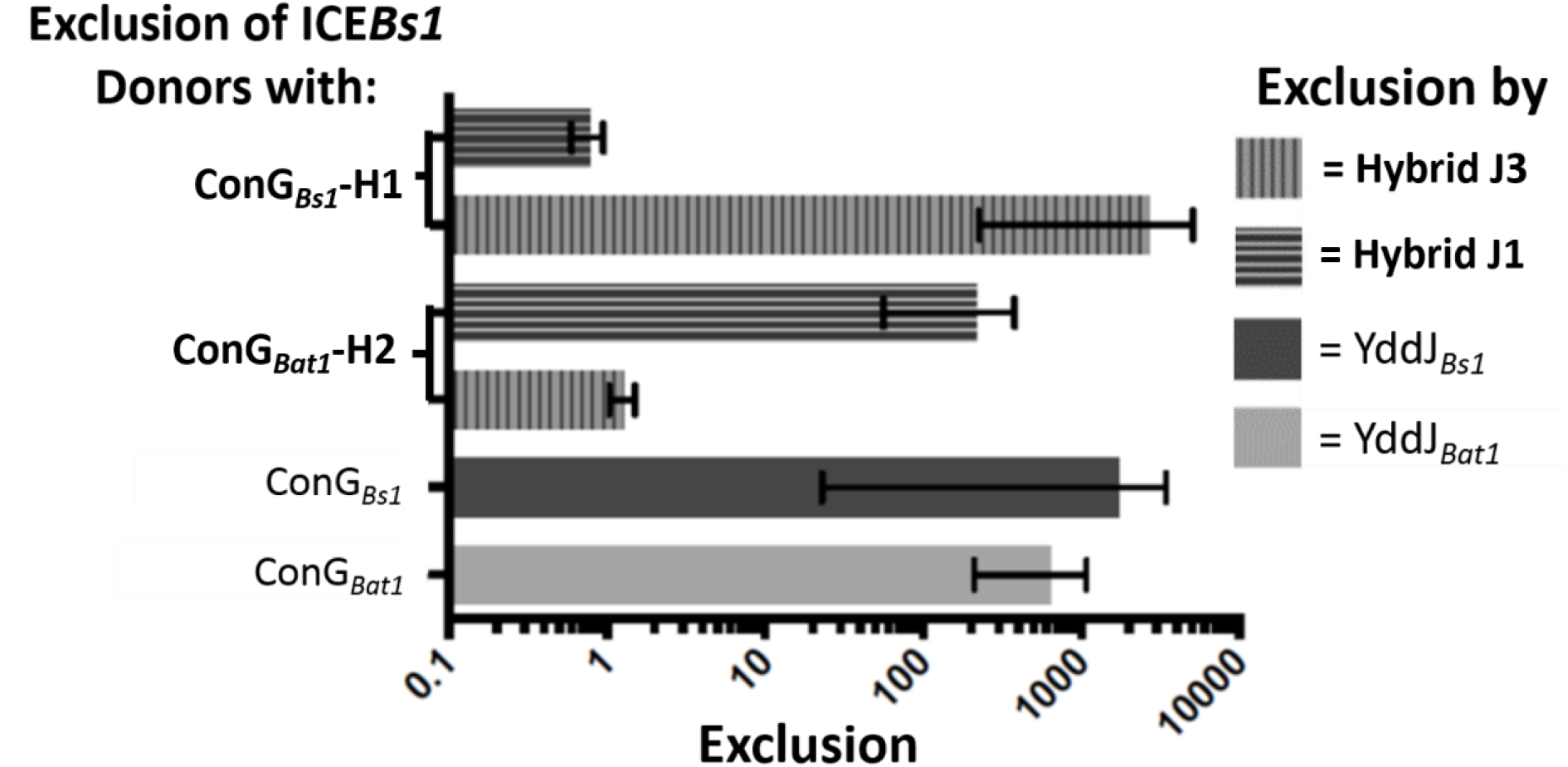
YddJ hybrid proteins can exclude conjugation machinery with ConG hybrid proteins. Conjugation machinery in the donor contained ConG_Bs1_-H1 (KPD136); ConG_Bat1_-H2 (KPD135); ConG_Bs1_ (KPD225); ConG_Bat1_ (KPD224). Recipients expressed YddJ_Bs1_ (black bars; MA982); YddJ_Bat1_ (gray bars; KPD219); hybrid J3 (vertical stripes; KPD128); hybrid J1 (horizontal stripes; KPD132). Exclusion was calculated as for Fig. 2. with results from matings into recipients that did not contain *yddJ* (CAL89) that were done in parallel. Data bars represent averages from three independent experiments, with error bars depicting standard deviations.

### Death of ICE*Bs1* donors due to excessive mating

Previous work found that loss of exclusion leads to a drop in cell viability under conditions that support mating (12). However, it is not known if the drop in viability was due to killing of donors, recipients, or both. Experiments described below demonstrate that decreased viability of exclusion-defective mutants occurs when cells function concurrently as both donors and recipients.

There was considerable death of ICE*Bs1* host cells (initial donors) when these cells were surrounded by an excess of recipient cells. We mixed ICE*Bs1* donors that had exclusion (KPD154) with recipient cells at a ratio of 1 donor to ~100 recipients. After mating on filters, there was a dramatic drop in viability of the original donor cells such that only ~5% (4.6 ± 2.0%) of the original donors survived post-mating (Fig. 4). In the absence of exclusion (*ΔyddJ*, strain KPD155), only ~2% (1.8 ± 0.5%) of the original donors survived (Fig. 4), a significant difference based on the one-tailed t test (p = 0.0174).

**Figure 4.**
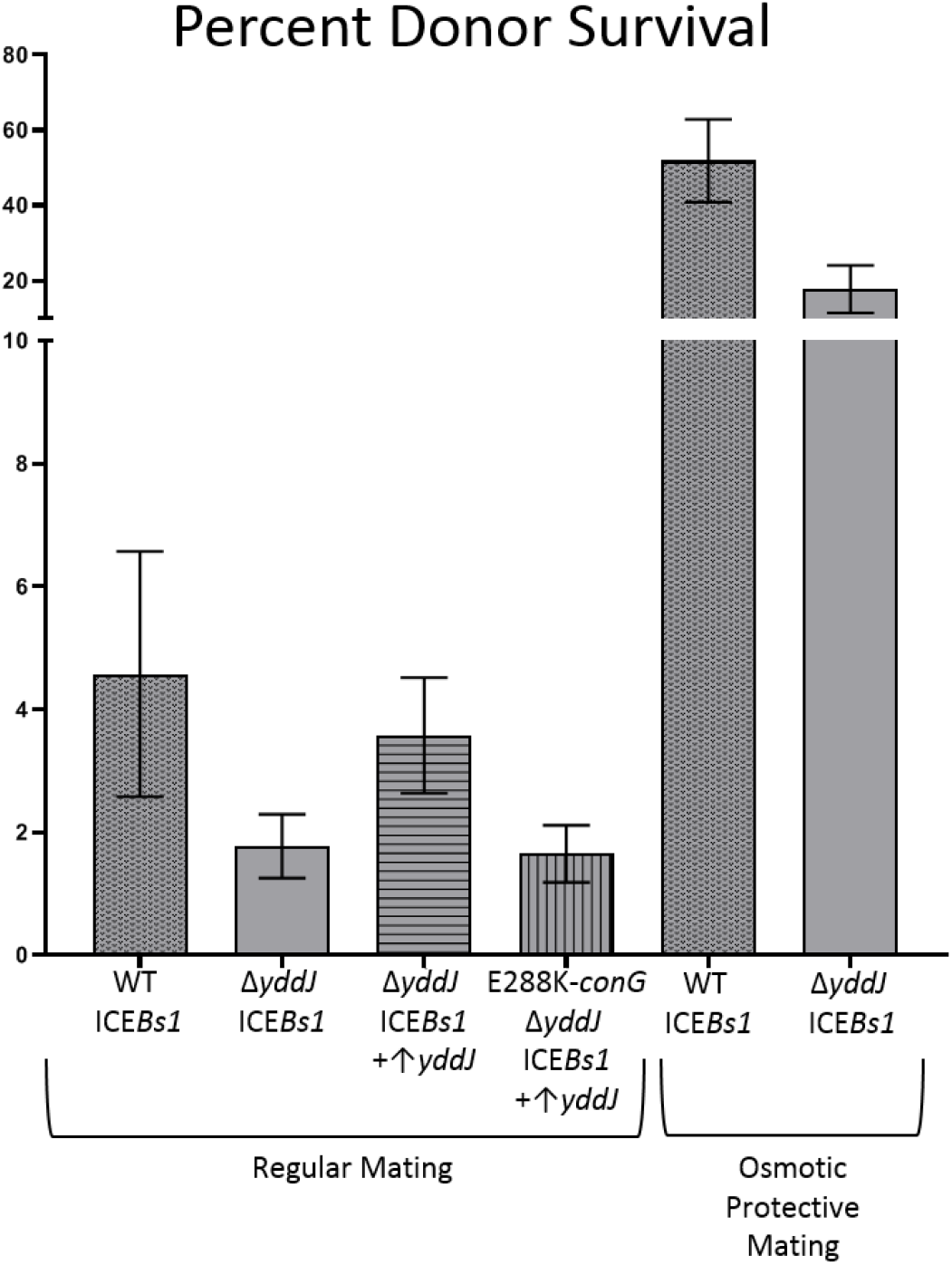
Death of donors by excessive mating is exacerbated by loss of exclusion and largely alleviated under osmo-protective conditions. WT ICE*Bs1* donors were mixed with recipients that lacked ICE (CAL419) at a ratio of 1 donor to ~100 recipients and put through mating conditions described in Fig. 1. In this case, mating filters were placed on agar plates as above (regular mating) or with osmo-protection. After incubation, mating mixtures were resuspended either without (regular mating) or with osmo-protection. Percent donor survival was determined by measuring CFUs/ml after mating compared to that prior to mating. Donors included: ICE*Bs1* (wt; KPD154); ICE*Bs1* Δ*yddJ* (KPD155); ICE*Bs1* Δ*yddJ* overexpressing *yddJ* from an ectopic locus (KPD156); ICE*Bs1* Δ*yddJ conG*E288K (resistant to exclusion), also overexpressing *yddJ* from an ectopic locus (KPD157). Data bars represent averages from the three replicate mating assays for each donor, with error bars depicting standard deviations. p-values from one-tailed t test were: 0.0174 for wild type compared to *ΔyddJ*; 0.0052 for *ΔyddJ* with over-expressed *yddJ* compared to *conGE288K* (exclusion-resistant) with over-expressed *yddJ*; and 8.43 × 10^−4^ for the pair compared under osmo-protective conditions.

The decrease in donor survival was due to loss of exclusion and not the absence of YddJ per se. We analyzed survival of donors that express YddJ, but that contain a missense mutation in *conG* that make them insensitive to exclusion (12). This mutant also had decreased survival (1.7 ± 0.5%), similar to that of cells without *yddJ* (Fig. 4). Together, these results indicate that 1) there is significant donor death when the donors are surrounded by a vast excess of recipients and likely to be transferring to multiple cells; and 2) that exclusion provides ~2-3-fold protection from this death.

The protection conferred by YddJ was due to its function in the original donors. We expressed *yddJ* from an ectopic locus in donor cells that contained ICE*Bs1ΔyddJ* (KPD156). The original donor has exclusion, but a transconjugant will not because the transferred element (ICE*Bs1ΔyddJ*) is missing *yddJ*. Survival of donors ectopically expressing *yddJ* was ~4% (3.6 ± 0.9%) (Fig. 4) in mating experiments analogous to those above with 1 donor to ~100 recipients. These results indicate that in the absence of exclusion, some of the donor death is likely due to donors acting as recipients in conjugation and that the transconjugants are likely transferring DNA back to the original donors. In donors capable of excluding entry of a second copy of ICE*Bs1*, donor death is likely from mating events with many recipients.

The drop in viability of donor cells was due to the presence of the mating machinery and close proximity of recipients. Donor death was dependent on overexpression of *rapI* to induce ICE*Bs1*. Without *rapI* induction, there was no detectable drop in donor cell viability under conditions that mimic the mating described above. Furthermore, donor death was not simply due to overexpression of *rapI* and activation of ICE*Bs1*. Matings done with both wild type ICE*Bs1* donors (KPD154) and Δ*yddJ* ICE*Bs1* donors (KPD155) at a ratio of 1 donor to ~100 recipients, with overexpression of *rapI*, but at a cell concentration low enough to reduce mating (~4×10^5 rather than ~8×10^8 total cells for mating) had 63 ± 16% and 89 ± 7% survival of wild type and *yddJ* donors, respectively. Thus, the large drop in viability was dependent on activation of ICE*Bs1* and conditions that support multiple mating events. We postulate that excessive mating, and serving as both a donor and recipient (in the absence of YddJ-mediated exclusion), likely causes cell wall damage that leads to cell death.

### Osmo-protective conditions increase survival during excessive mating

We found that osmo-protective conditions increased donor survival under conditions of excessive mating, both with and without exclusion. Mating assays were done using donors with (KPD154) and without (ICE*Bs1ΔyddJ*, strain KPD155) exclusion at a ratio of 1 donor to ~100 recipients (CAL419) in osmo-protective conditions. These osmo-protective conditions consisted of replacing 1X Spizizen’s salts used as the support for mating filters and used to recover cells from the mating filters, with a solution of 20mM MgCl2 and 0.5M sucrose and buffered with 20mM maleic acid (see Materials and Methods). This solution (called MSM) has been used for the propagation *B. subtilis* cells with no cell wall, so-called L-forms (28).

In matings with osmo-protective conditions, survival of wild type donors was 51.9 ± 11.0%, and survival of donors without exclusion (ICE*Bs1*Δ*yddJ*) was 17.7 ± 6.3%, compared to 4.6 ± 2.0% and 1.8 ± 0.5%, respectively, in matings without osmo-protection (Fig. 4). Thus, a significant amount of donor death for both wild type and exclusion-deficient donors was eliminated by osmo-protection. These results indicate that donors surrounded by an excess of recipients are likely dying from cell wall damage due to excessive mating, either into a single recipient or into multiple recipients. There was still lower survival of donors without exclusion, even under the osmo-protective conditions used here. We suspect that this is due to the shift from osmo-protection to LB agar plates, and that the donors without exclusion have more envelope damage and lower survival following the shift.

## Discussion

Results presented here show that *B. subtilis* cells that transfer ICE*Bs1* can die from excessive transfer. This death is exacerbated by loss of exclusion, which likely enables transfer from a transconjugant back into the original donor. Death of the donors is largely relieved under osmo-protective conditions, indicating that death is due to alterations in the integrity of the cell envelope. If there is death of recipients we would not have detected it in our assays.

### Death of ICE*Bs1* donors compared to lethal zygosis in *E. coli*

Death of ICE*Bs1* donors and the protection by exclusion is different from the previously characterized phenomenon of lethal zygosis in *E. coli*. In lethal zygosis, cells are killed when they serve as recipients during multiple conjugation events. This killing occurs when recipients lacking the F plasmid (F-) are mixed with an excess of either Hfr donors or F+ exclusion-deficient donors. Recipient death by lethal zygosis also occurs when F+ exclusion-deficient recipients are mixed with an excess of Hfr donors (15–18). This killing is probably caused by increased permeability of the cell wall due to multiple matings (15). During Hfr transfer, the transconjugants do not become donors because the entire conjugative element is not transferred. This is in contrast to the situation with ICE*Bs1* in which transconjugants acquire the entire element and quickly become donors (29). It is not known if *E. coli* donors also die under conditions of excess mating.

### Benefits of ICE*Bs1* exclusion

The protective benefit of ICE*Bs1* exclusion probably serves an important role when ICE*Bs1* is breaking into a new population of host cells, a situation that is likely mimicked by matings with 1 donor to ~100 recipients. Once a cell receives ICE, it is ready to quickly donate it to other cells, and this feature gives ICE*Bs1* some distinct advantages, like being able to move quickly through cell chains via conjugation (29) and spread in a biofilm (30,31). That transconjugants quickly become donors indicates that if a mating pair is reasonably stable, then there is the likelihood that a transconjugant could transfer ICE back to the original donor. Our results indicate that it is taxing for ICE*Bs1* host cells to serve as donors. There is a considerable amount of donor death with multiple transfer opportunities (1 donor per 100 recipients at high cell concentrations), and there is more death in the absence of exclusion. Whatever the mechanism of killing, it seems that exclusion can protect ICE*Bs1* donor cells when they are already in the vulnerable state of serving or having just served as donors.

### Comparison of exclusion proteins

Exclusion proteins from different families of conjugative elements have limited sequence similarity, but still have some common features. In general, exclusion proteins are relatively small and found on the surface of the host cell where it is in position to inhibit cognate conjugation machinery in a potential donor (14). Exclusion proteins are usually not required for conjugative transfer, except for that of R27 (32), often function in a dose-dependent manner, and target a VirB6 (ConG) homolog or analog in the cognate secretion system, as discussed above.

In the case of the F/R100 family of plasmids (*E. coli, S. flexneri*), the exclusion protein TraS a small hydrophobic protein except for a short hydrophilic region (33), predicted to be localized to the inner membrane (23). In the SXT/R391 family of ICEs (*V. cholerae, P. rettgeri*), the exclusion protein Eex is in the inner membrane (26), and paradoxically, the regions essential for exclusion specificity are in a cytoplasm region of the protein (34). For ICE*Bs1* (*B. subtilis*), the exclusion protein YddJ is predicted to be extracellular and attached to the cell surface via a lipid modification (12).

### Targets of exclusion proteins

The targets of exclusion proteins from four different families of conjugative elements, all from gram-negative bacteria, have been identified. In each case, the exclusion protein targets its cognate VirB6 homologue or analogue. The region conferring specificity of exclusion appears to be either periplasmic (23) or cytoplasmic (24,26,34). Likewise, the region of ConG of ICE*Bs1* (and ICE*Bat1*) that confers exclusion specificity is predicted to be cytoplasmic. Together, these analyses indicate that either this normally periplasmic or cytoplasmic region can be present on the cell surface, or that the cognate exclusion protein has access to part of the periplasm or cytoplasm.

### Specificity of exclusion and contributions to ICE biology

Identification of key regions for exclusion specificity in ConG and YddJ also highlight important aspects of ICE*Bs1* biology, and how ICEs contribute to bacterial evolution by spreading genetic material. The fact that exclusion (or lack thereof) can be based on differences of a few residues in the exclusion protein or target protein demonstrates that exclusion by a copy of ICE*Bs1* can very selectively allow slightly different elements (such as ICE*Bat1*) to enter the host cell, while significantly reducing the number the number conjugation attempts by other would-be ICE*Bs1* donors.

ICEs play an important role in bacterial evolution by contributing to the spread of genetic material, and one way in which an ICE gains or loses genetic material (which it can then transfer along with itself) is through genetic rearrangement events with other ICEs and plasmids (5). It has been theorized (14) that the lack of exclusion systems in some ICEs allows for more rapid evolution of the ICE, but this could be harmful for ICE*Bs1* given its strict requirement for integration site. Having an exclusion system that allows for as much exposure to other elements as possible, while limiting the number of identical elements that enter, would allow ICE*Bs1* to have the chance to be exposed to as many other ICEs as possible and benefit from the genetic diversity, while avoiding suffering the ill effects.

## Materials and Methods

### Media and growth conditions

Cells were typically grown in S7_50_ defined medium (35) supplemented as needed for auxotrophic requirements (40 μg/ml tryptophan, 40 μg/ml phenylalanine, and 200 μg/ml threonine for strains containing alleles inserted at *thrC*). Isopropyl-β-D-thiogalactopyranoside (IPTG, Sigma) was used at a final concentration of 1 mM to induce expression from the promoter Pspank(hy). LB plates contained antibiotics, where indicated, at the following concentrations: kanamycin (5 μg/ml), spectinomycin (100 μg/ml), streptomycin (100 μg/ml), and a combination of erythromycin (0.5 μg/ml) and lincomycin (12.5 μg/ml) to select for macrolide-lincosamide-streptogramin (MLS) resistance.

### Strains and alleles

*B. subtilis* strains used in this study are listed in Table 1. Cloning and generation of strains was done following standard techniques (36). All strains (KPD219, CAL89, MA982, KPD128, KPD131, KPD132, KPD137, CAL419) used as recipients in mating experiments did not contain ICE*Bs1* (ICE^0^), contained null mutations in *comK* or *comC* (described below), and were streptomycin-resistant (*str-84*) (9,37). Streptomycin was used as a counter-selective marker in mating assays (see more on mating assays below).

**Table 1.**
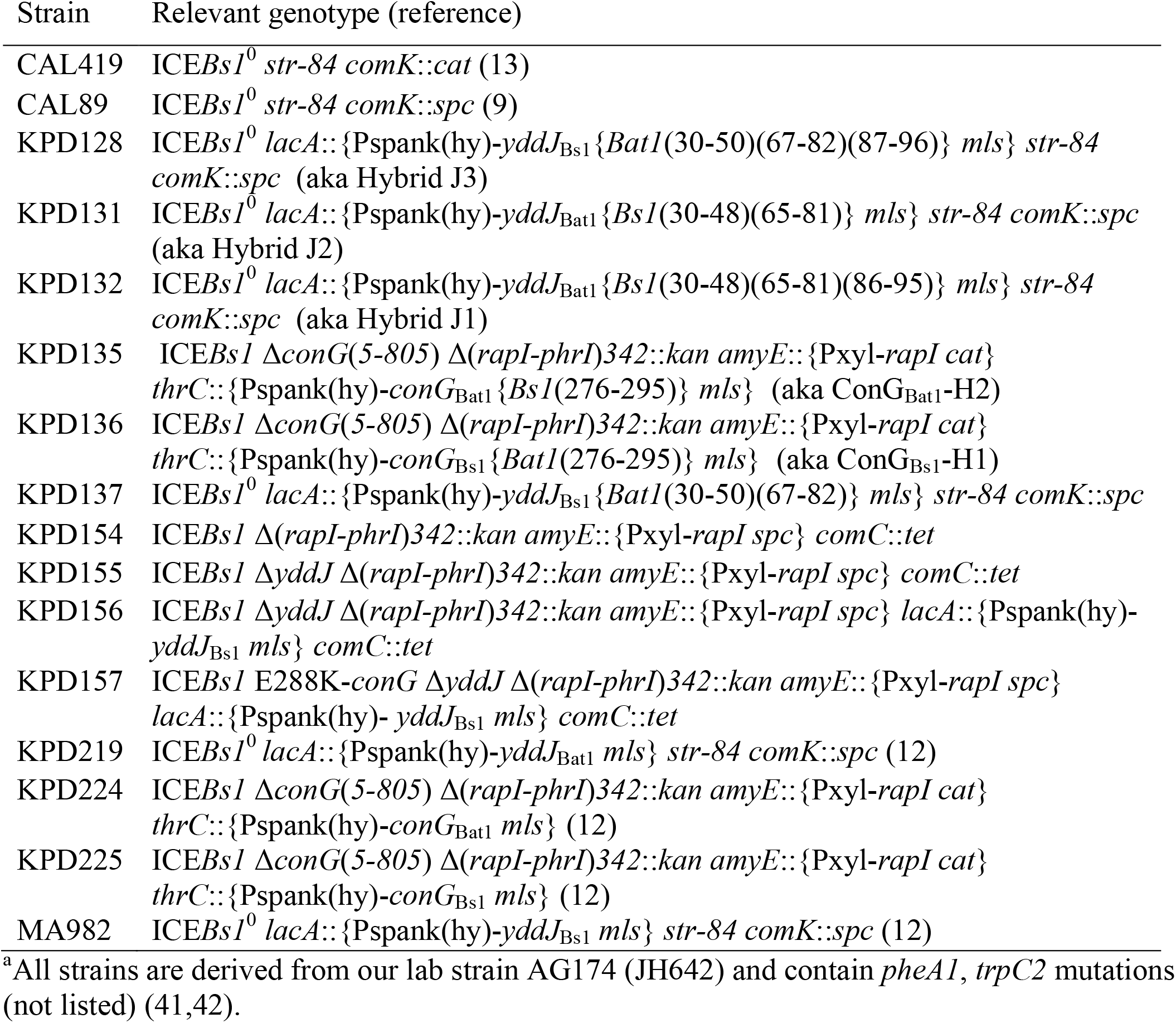
*B. subtilis* strains used^a^

All ICE*Bs1* donor strains contain a version of ICE*Bs1* that has a kanamycin-resistance gene inserted in place of *rapI-phrI*: Δ(*rapI*-*phrI*)*342*::*kan* (9). *rapI* was over-expressed from Pxyl-*rapI* in donor cells, to achieve inducible ICE*Bs1* gene expression and excision. Pxyl-*rapI* alleles were integrated into *amyE* with *spc* or *cat* antibiotic resistance genes: *amyE*::{Pxyl-*rapI spc*} (38) or *amyE*::{Pxyl-*rapI cat*} (38,39). Any ICE*Bs1* donor strains containing a deletion of *conG*, Δ*conG*(*5-805*), were derived from MMB1283 (27). KPD210, a donor strain containing a complete deletion of *yddJ*, was derived from MA11 (12).

#### Construction of *comK* and *comC* null mutations

Null mutations in *comK* and *comC* were used to prevent transformation in all recipient strains, and in donor strains (KPD154, KPD155, KPD156, KPD157) used in experiments where even low levels of transformation could significantly alter donor CFU counts. *comK*::*cat* was from CAL419 (37), *comK*::*spc* (9) and *comC::mls* (12) were also previously described.

#### Construction of Pspank(hy)-*yddJ* and Pspank(hy)-*yddJ* chimeras at *lacA*

All *yddJ* overexpression constructs consist of *yddJ* fused to the LacI-repressible IPTG-inducible promoter Pspank(hy), and integrated at *lacA* with an *mls* resistance gene. Pspank(hy)-*yddJ*_Bs1_ (*yddJ* from ICE*Bs1*) present in strain MA982, and Pspank(hy)-*yddJ*_Bat1_, (*yddJ* from ICE*Bat1*) present in strain KPD219, were described previously (12).

To make the *yddJ* hybrids, *yddJ*_Bat1_ DNA was amplified by PCR from genomic DNA from *B. atrophaeus* strain 11A1 (from the Bacillus Genetic Stock Center; www.bgsc.org) and *yddJ*_Bs1_ DNA was amplified by PCR from genomic DNA from *B. subtilis* strain AG174. Fragments of each *yddJ* were amplified and fused by isothermal assembly as necessary to make four chimeric constructs.

1. Hybrid J1: formally known as YddJ_Bat1_{*Bs1*(30-48)(65-81)(86-95)}, in which YddJ_Bs1_ residues 30-48, 65-81, and 86-95 were substituted for their corresponding YddJ_Bat1_ residues (30-50, 67-82, and 87-96, respectively).
2. Hybrid J2: formally known as YddJ_Bat1_{*Bs1*(30-48)(65-81)}, in which YddJ_Bs1_ residues 30-48 and 65-81 were substituted for their corresponding YddJ_Bat1_ residues (30-50 and 67-82, respectively).
3. Hybrid J3: formally known as YddJ_Bs1_{*Bat1*(30-50)(67-82)(87-96)}, in which YddJ_Bat1_ amino acids 30-50, 67-82, and 87-96 were substituted for their corresponding YddJ_Bs1_ amino acids (30-48, 65-81, and 86-95, respectively).
4. Hybrid J4: formally known as YddJ_Bs1_{*Bat1*(30-50)(67-82)}, in which YddJ_Bat1_ residues 30-50 and 67-82 were substituted for residues 30-48 and 65-81, respectively, in YddJ_Bs1_.

For all Pspank(hy)-*yddJ* constructs, the *yddJ* PCR fragments were joined together, and then joined to two fragments amplified from pCJ80 (a vector for making fusions to Pspank(hy) and integration at *lacA*) (40) by isothermal assembly. One fragment from pCJ80 included the pCJ80 SphI cut site and 2409 bp upstream of this cut site, including homology to the 5’ end of *lacA*. The other fragment included the pCJ80 SacI cut site and 2299 bp downstream of this cut site, including homology to the 3’ end of *lacA*. These two fragments were digested with SphI and SacI, respectively, before isothermal assembly with the *yddJ* PCR DNA. The primers used to amplify *yddJ* contained sequences complementary to sequences in the primers used to amplify regions of *lacA*, thereby enabling joining by isothermal assembly. The resulting isothermal assembly product was integrated by double cross-over into the chromosome by transformation and selecting for MLS resistance, to generate the *yddJ* overexpression alleles.

#### Construction of Pspank(hy)-*conG* and Pspank(hy)-*conG* chimeras at *thrC*

*conG* was expressed ectopically from the LacI-repressible IPTG-inducible promoter Pspank(hy), from constructs integrated at *thrC* with an *mls* resistance gene. The Pspank(hy)-*conG* alleles were used to complement the Δ*conG*(*5-805*) deletion in ICE*Bs1*. Pspank(hy)-*conG*_Bs1_ (*conG* from ICE*Bs1*) present in strain KPD225, and Pspank(hy)-*conG*_Bat1_, (*conG* from ICE*Bat1*) present in strain KPD224, have been described (12,27).

To make the *conG* hybrids, *conG*_Bat1_ DNA was amplified by PCR from genomic DNA from *B. atrophaeus* strain 11A1 (from the Bacillus Genetic Stock Center; www.bgsc.org) and *conG*_Bs1_DNA was amplified by PCR from genomic DNA from *B. subtilis* strain AG174. Fragments of each *conG* were amplified and fused by isothermal assembly as necessary to make two chimeric constructs.

1. ConG_Bs1_-H1: formally known as Con*G*_Bs1_{*Bat1*(276-295)}, in which ConG_Bat1_ residues 276-295 were substituted for residues 276-295 in ConG_Bs1_.
2. ConG_Bat1_-H2: formally known as ConG_Bat1_{*Bs1*(276-295)}, in which ConG_Bs1_ residues 276-295 were substituted for residues 276-295 in ConG_Bat1_.

For all Pspank(hy)-*conG* constructs, the *conG* PCR fragments were joined together, and then joined to two fragments amplified from pMMB1341 (27) by isothermal assembly. One fragment from pMMB1341 included the HindIII cut site and the adjacent 2330 bp upstream of this cut site, which includes sequences from the 3’ end of *thrC*. The other fragment included the SphI cut site and the adjacent 1867 bp downstream, which includes sequences from the 5’ end of *thrC*. These two fragments were digested with HindIII and SphI, respectively, before isothermal assembly with the *conG* PCR DNA. The primers used to amplify *conG* contained sequences complementary to sequences in the primers used to amplify regions of *thrC*, thereby enabling joining by isothermal assembly. The resulting isothermal assembly product was integrated by double cross-over into the chromosome by transformation and selecting for MLS resistance, to generate the *conG* overexpression alleles.

### Mating Assays

Mating assays were performed essentially as described (9,37). Donor and recipient cultures were grown in S7_50_ defined minimal medium supplemented with 0.1 % glutamate and 1% arabinose until they reached mid-exponential growth phase, then diluted back to an optical density at 600 nm (OD600) of 0.1. At this point 1% xylose was added to donor cultures to induce expression of Pxyl-*rapI*. After two hours of xylose induction, donor and recipient cells were mixed, and poured over a nitrocellulose filter under vacuum filtration. Unless otherwise indicated, equal numbers of donor and recipient cells were used (~4×10^8^ cells of each). Filters were incubated for 3 hours at 37°C on 1.5% agar plates containing 1x Spizizen’s salts (2 g/l (NH_4_)SO_4_, 14 g/l K_2_HPO_4_, 6 g/l KH_2_PO_4_, 1 g/l Na_3_ citrate-2H_2_O, 0.2 g/l MgSO_4_-7H_2_0) (36). Cells were re-suspended from the filters, serially diluted in 1X Spizizen’s salts and plated on LB agar plates containing kanamycin and streptomycin to select for transconjugants. For matings done in osmo-protective conditions, 1X Spizizen’s salts (in mating plates and resuspension media) was replaced with MSM (20mM MgCl2, 0.5M sucrose and buffered with 20mM maleic acid) (28). The number of viable ICE*Bs1* donor cells (CFU/ml) was determined at the time of donor and recipient cell mixing, by serial dilution plating on LB agar plates containing kanamycin. Mating efficiency was calculated as the percent transconjugants CFU/ml per donor CFU/ml (at the time of mixing donors with recipients). The fold-exclusion was calculated as the percent transfer into an ICE^0^ recipient divided by the percent transfer into an ICE^0^ recipient that was expressing *yddJ*.

## Acknowledgements

We thank Janet Smith for helpful comments on the manuscript. This research was supported, in part, by the National Institute of General Medical Sciences of the National Institutes of Health under award number R01 GM050895 and R35 GM122538 to ADG. KPD was also supported, in part, by the NIGMS predoctoral training grant T32 GM007287. Any opinions, findings, and conclusions or recommendations expressed in this report are those of the authors and do not necessarily reflect the views of the National Institutes of Health.

